# Spatial control of gene expression in flies using bacterially derived binary transactivation systems

**DOI:** 10.1101/2020.11.24.396325

**Authors:** Stephanie Gamez, Luis C. Vesga, Stelia C. Mendez-Sanchez, Omar S. Akbari

## Abstract

Controlling gene expression is an instrumental tool for biotechnology, as it enables the dissection of gene function, affording precise spatial-temporal resolution. To generate this control, binary transactivational systems have been used employing a modular activator consisting of a DNA binding domain(s) fused to activation domain(s). For fly genetics, many binary transactivational systems have been exploited *in vivo*; however as the study of complex problems often requires multiple systems that can be used in parallel, there is a need to identify additional bipartite genetic systems. To expand this molecular genetic toolbox, we tested multiple bacterially-derived binary transactivational systems in *Drosophila melanogaster* including the *p*-CymR operon from *Pseudomonas putida*, PipR operon from *Streptomyces coelicolor*, TtgR operon from *Pseudomonas putida*, and the VanR operon from *Caulobacter crescentus*. Our work provides the first characterization of these systems in an animal model *in vivo*. For each system we demonstrate robust tissue-specific spatial transactivation of reporter gene expression, enabling future studies to exploit these transactivational systems for molecular genetic studies.

## Introduction

Precise regulation of gene expression is instrumental in biological applications such as therapeutics (Kemmer et al. 2010) and pharmaceuticals (Sharpless and Depinho 2006), where long-term regulation of gene expression for gene therapy is crucial following rational cell reprogramming in tissue engineering (Fussenegger et al. 1998) or is required to build sensors for synthetic gene circuits (Deans, Cantor, and Collins 2007; Kramer and Fussenegger 2005). This precise control is currently afforded by synthetic binary expression systems, which are engineered control systems that can oftentimes respond to the presence of modified proteins and chemical molecules (ligands). More specifically, these systems generally couple a synthetic transcription factor with a transactivation domain that binds to specific operator sites (Triezenberg, Kingsbury, and McKnight 1988). These systems can control gene expression in a temporal- and tissue-specific manner, using appropriate regulatory elements to express transactivators. This controlled expression is especially important for toxic gene products or temporal/tissue-specific knock-down of an essential gene, which would be otherwise impossible due to their deleterious effect on the organism.

While several binary transactivation systems exist, only a handful have been shown to function *in vivo* in *Drosophila melanogaster* (reviewed (Venken, Simpson, and Bellen 2011)), therefore we sought to further expand this powerful molecular genetic toolbox. For example in flies, transactivation systems have been used extensively *in vivo* affording spatial control including: Gal4-UAS adapted from yeast (Brand and Perrimon 1993), the Q-system adapted from the bread mold *Neurospora crassa (Potter et al. 2010)*, and several systems derived from bacteria including the LexA/LexAop (Lai and Lee 2006), the Tet system using tTA/TRE (Bello, Resendez-Perez, and Gehring 1998), transcription activator-like effectors (TALEs)(Toegel et al. 2017), and recently even CRISPR/dCas9-VPR based transactivators (Lin et al. 2015; Jia et al. 2018). In addition to spatial control, some of these systems also afford temporal control by exploiting small-molecule triggers to fine-tune expression in a dose-dependent manner. For example, the Gal4-UAS system utilizes mifepristone (RU486) (Nicholson et al. 2008; Robles-Murguia et al. 2019), and temperature by using a temperature sensitive allele of GAL80, GAL80ts (McGuire et al. 2003), or trimethoprim by incorporating a destabilizing domain (Sethi and Wang 2017). Other chemically-controlled systems include the Tet system which uses tetracycline/doxycyline (Bello, Resendez-Perez, and Gehring 1998; Bieschke, Wheeler, and Tower 1998), and the Q-system which uses quinic acid (Potter et al. 2010).

Despite this desirable level of precise spatial-temporal control, concerns have been raised over their potential side-effects in animals. For example, due to the negative fitness effects of RU486 at certain concentrations, the Gal4-UAS system may not be ideal (Landis et al. 2015; Yamada et al. 2017). Moreover, the use of temperature in flies can have a significant impact on the behavior and physiology (Parisky et al. 2016). Tetracycline/doxycycline has also been reported to have negative physiological impacts (Chatzispyrou et al. 2015; Moullan et al. 2015), including impaired mitochondrial function (Zeh et al. 2012), which may affect experimental outcomes. While the Q-system has been demonstrated to be efficient and has no documented side effects using quinic acid, this system requires both an additional genetic component, termed QS, to suppress gene expression and the supplementation of quinic acid for de-repression of QS protein (Potter et al. 2010).

Herein, we sought to characterize additional binary systems to expand the *Drosophila* molecular genetic tool box. We tested four bacterially derived systems by encoding them in *Drosophila* strains including, *p*-CymR operon from *Pseudomonas putida* (Mullick et al. 2006), PipR operon from *Streptomyces coelicolor (Fussenegger et al. 2000)*, TtgR operon from *Pseudomonas putida* (Gitzinger et al. 2009), and the VanR operon from *Caulobacter crescentus* (Gitzinger et al. 2012). To characterize these systems, we exploited a novel dual-luciferase reporter system incorporating GFP enabling both quantification and visualization of gene-expression levels, respectively, as compared to the widely used Tet-OFF system (tTA). Additionally, we tested the systems’ ability to be controlled via small molecules, which may provide avenues for further optimization. Overall, our results demonstrate the robust spatial transactivational potential of these control reporter systems and validate their relevance for future studies. This work is the first report of these particular prokaryotic systems engineered in *Drosophila* and provides the field with additional spatially-controlled transactivational systems that can be used as genetic circuits.

## Results

### Design and development of additional binary systems in flies

To characterize the utility of bacterially derived transactivation systems in *D. melanogaster*, including: the *p*-CymR operon from *Pseudomonas putida* (Mullick et al. 2006), the PipR operon from *Streptomyces coelicolor (Fussenegger et al. 2000)*, the TtgR operon from *Pseudomonas putida* (Gitzinger et al. 2009), the VanR operon from *Caulobacter crescentus* (Gitzinger et al. 2012), we designed a dual luciferase reporter system utilizing the repressor and corresponding operator sequences from each bacterial operon. The widely used TetR system served as a positive control (Bello, Resendez-Perez, and Gehring 1998). Separate “driver” and “responder” transgenic lines were generated that could be genetically crossed to visualize and quantify transactivation responses (**Fig. 1A, Fig. S1A**). In each driver line, a *flightin* (*Fln*) promoter fragment (Ayer and Vigoreaux 2003) was utilized to drive expression of a chimeric transactivator in the indirect flight muscles consisting of the operon specific repressor protein (i.e. CymR, PipR, TtgR, VanR, or TetR) fused to three tandem VP16 activation domains (Das, Tenenbaum, and Berkhout 2016; Wysocka and Herr 2003)(**Fig. 1B**, **Fig. S1A**). Each responder line was designed with 2-3 operator sequences, specific to each operon (i.e. CymO, PipO, TtgO, VanO, or TetO) and upstream of both a minimal Hsp70 (Amin et al. 1987) and a UASp (Rørth 1998) promoter driving expression of firefly luciferase reporter genes linked to a T2A-GFP marker to enable direct quantification and visualization of transactivation via luciferase and GFP, respectively. The construct was terminated by a baculovirus derived p10 3’ UTR known to increase efficiency of both polyadenylation and expression (Pfeiffer, Truman, and Rubin 2012) (**Fig. S1A**). Ubiquitously expressed renilla luciferase with a SV40 3’UTR was also added to the responder construct, oriented in the opposite direction, to enable the normalization of firefly luciferase expression from the same genomic context. Both the driver and responder constructs were marked with the mini-white transformation marker (Pirrotta 1988), and inserted using phiC31 site-specifically into the same 3^rd^ chromosomal site as the test system to enable direct comparisons (**Fig. 2A**). Transgenic flies were balanced and maintained as homozygous stocks.

**Figure 1.**
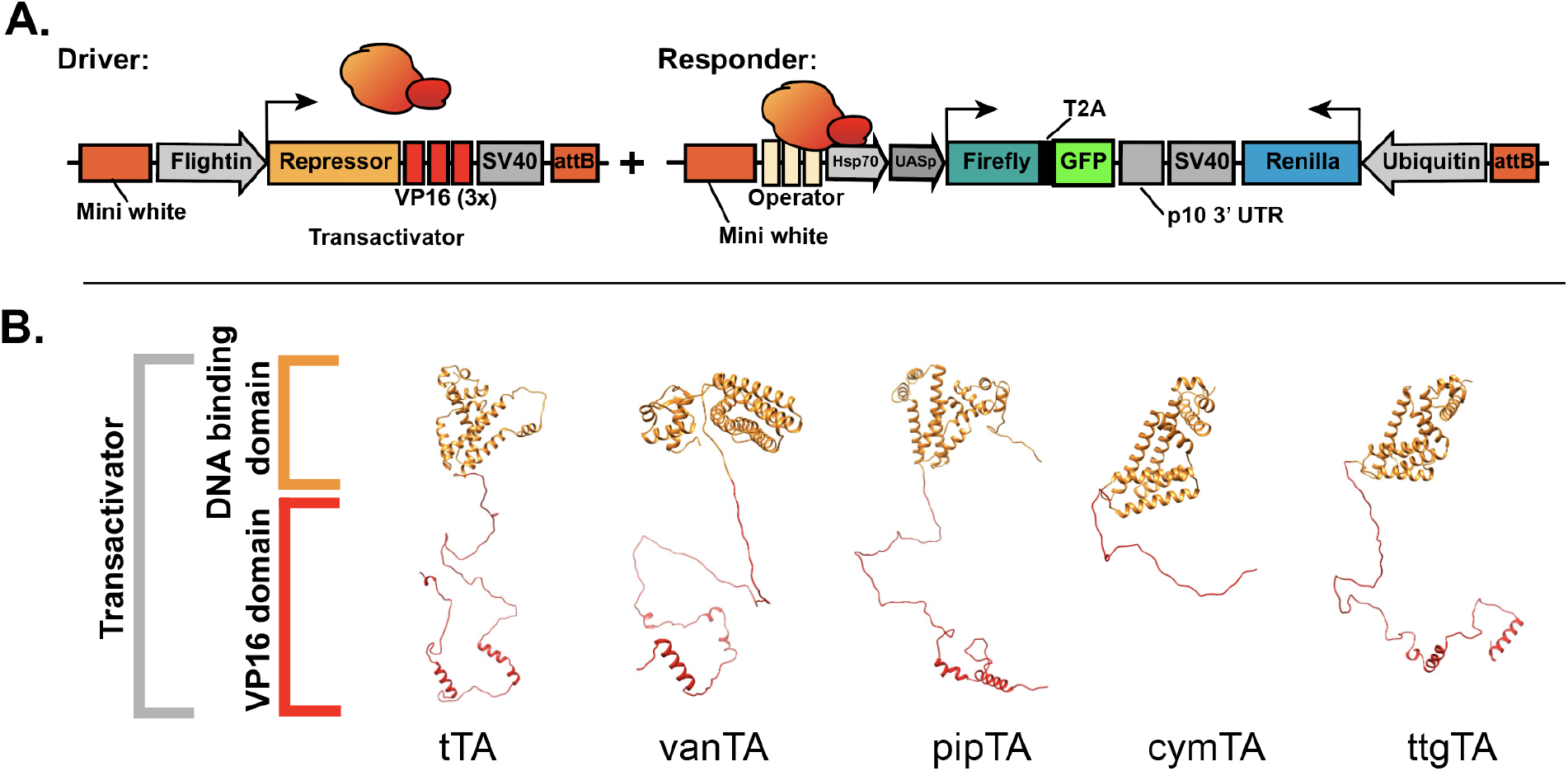
Characterization of transactivation systems. (A) Schematic of the general components in “driver” and “responder” transgenes. Driver transgenes are composed of a regulatory element driving the expression of a repressor fused to three tandem VP16 activation domains (transactivator). The responder transgene contains an operator sequence (2-3 copies), minimal promoters (HSP70 and UASp), firefly and renilla luciferase, and GFP to visualize and quantify transactivation. The transactivator (depicted in orange-red) binds to the operator sequence and induces the expression of reporter genes. A ubiquitous renilla luciferase in the responder transgene enables normalization of firefly luciferase. (B) 3D structural protein models (homology models) of system transactivators used in this study. Transactivators are made up of the DNA binding domain (shown in orange) and three tandem VP16 activation domains (in red). For each system, protein modeling was used to depict overall folding of fused transactivational components.

**Figure 2.**
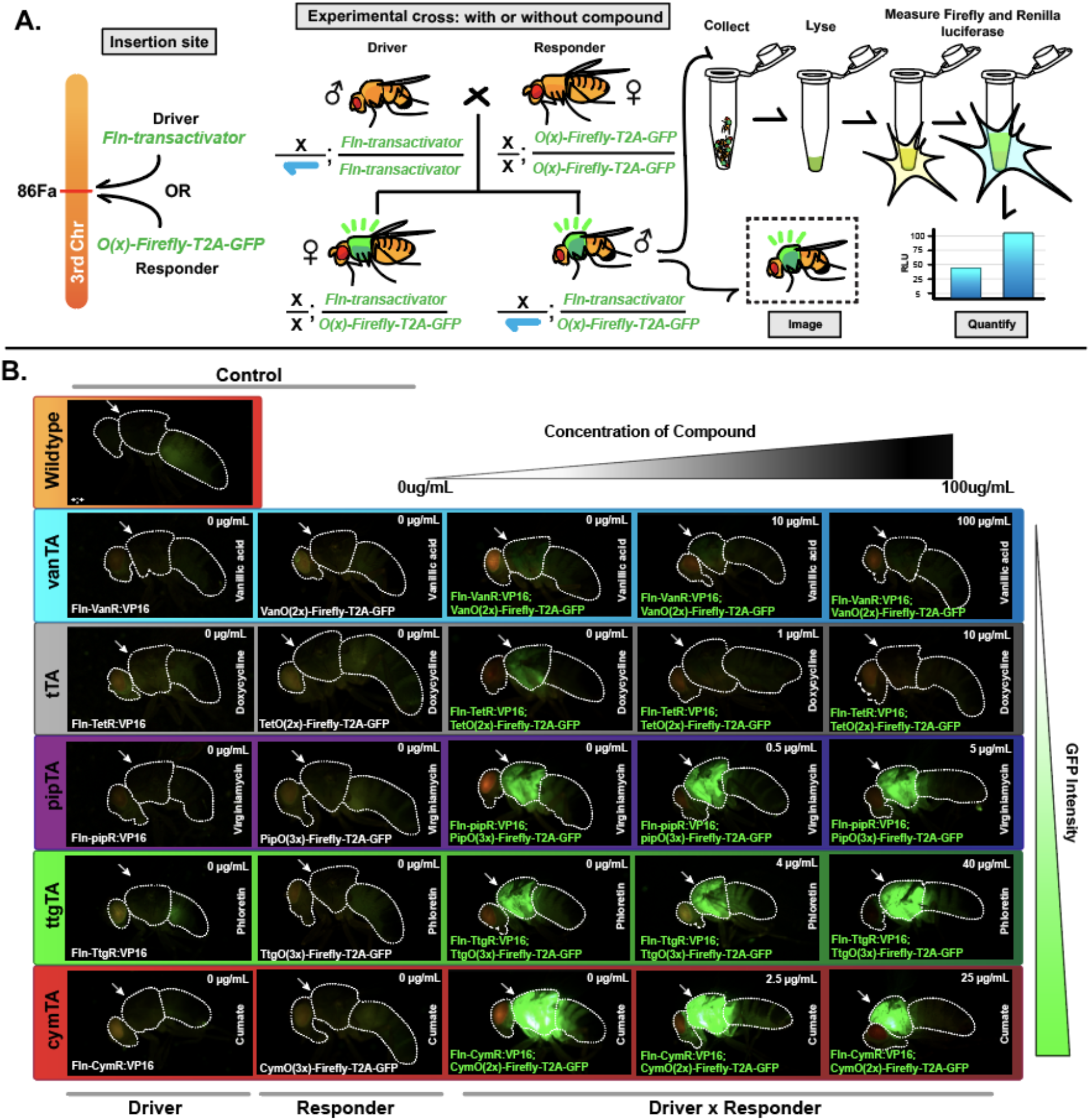
Characterization of alternative binary systems in *Drosophila melanogaster.* (A) Driver and responder transgenes were inserted site-specifically using a phiC31/attP docking site (ZH-attP-86Fa). A genetic cross between the homozygous driver and responder strains produced transheterozygous flies all displaying robust GFP fluorescence in the adult indirect flight muscles. Male flies were collected for imaging and quantification of dual luciferase reporters. (B) Transactivation of the GFP reporter was observed in all transheterozygotes in both the absence or presence of ligand. Robust GFP expression was observed in all transheterozygous (driver + responder) flies, while no GFP expression was observed in control files including wildtype, or driver-only, or responder-only files. White arrows point to the fly thorax/flight muscles where *Fln* is expressed. The ligand used is listed vertically on the right of each image. The ligand concentration fed to flies is indicated in the top right-hand corner of each image. Genotypes are shown in the bottom left corner. Only 1-2 day old males were imaged.

### Binary systems as transactivators of gene expression

To determine whether these bacterial systems were capable of transactivating reporter genes in flies, we first performed a genetic cross between the driver and responder lines to produce transheterozygotes (**Fig. 2A**). For each transhetrozygous transactivator/responder combination, robust GFP fluorescence was visible in the adult thorax where the flightN promoter was expressed in the indirect flight muscles, indicating that each combination was robustly transactivating (**Fig. 2B**). Importantly, aside from autofluorescence no basal GFP expression was observed in the control flies, which only harbored either the driver or responder, but not both. Differences in GFP fluorescence intensity across all systems, indicated system-specific differences in the levels of reporter gene expression, despite the fact that all the driver and responder transgenes were located on the same chromosomal site. Specifically, the CymR/CymO (cymTA) system had the highest visible GFP fluorescence, while the tetR/TetO (tTA) and the VanR/Van (vanTA) systems had the lowest (**Fig. 2B**). Because GFP fluorescence only provides a visual qualitative confirmation of transactivation, we next measured luciferase expression for an accurate quantification. To do this, both firefly and renilla luciferase were measured in individual 3–day-old transheterozygous flies using a dual luciferase assay (**Fig. 2A**). In all systems, transheterozygous flies had significant activation of firefly luciferase compared to control responder-only flies, suggesting robust transactivation for each system (**Fig. 3B**) (all systems had at least a *p* ≤ 0.005).

**Figure 3.**
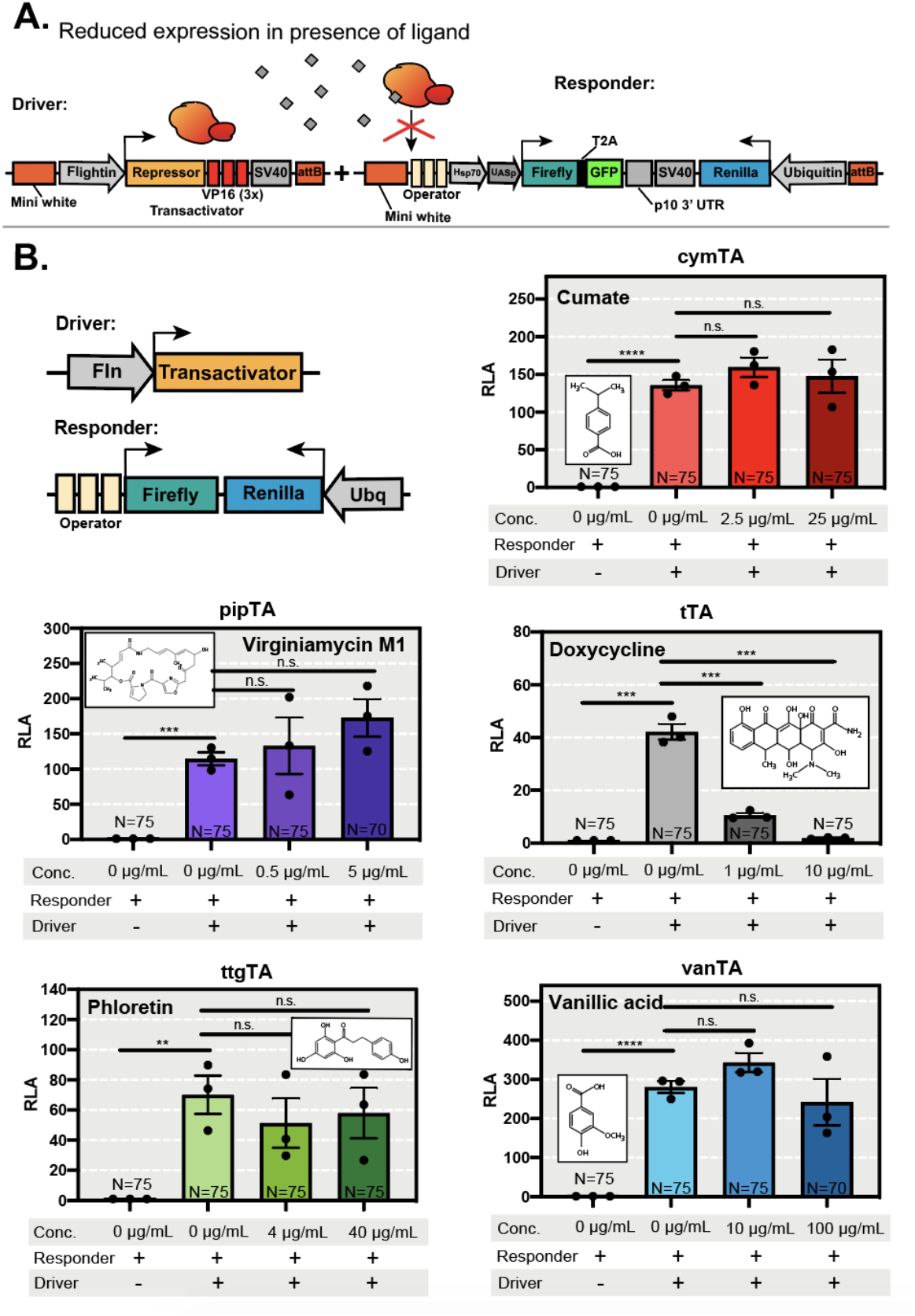
Transcriptional activity of transgenic flies with or without ligands. (A) We tested whether expression of reporter genes in the responder transgene depended on the presence or absence of ligand. In the absence of a ligand, the transactivator (depicted in orange-red) binds to the operator sequence and induces the expression of reporter genes. In the presence of a ligand (depicted as a gray diamond), we expected the ligand binding to the transactivator to prevent the transactivator from binding to the operator sequence, preventing reporter gene expression. (B) Relative luciferase activity (RLA) for responder-only flies (control) and transheterozygous flies on increasing concentrations of ligand. All systems show significant transactivation of firefly luciferase expression when both driver and responder transgenes are present in the same genomic context. Each system displays a unique expression level, depicting system-specific differences. When cymTA, pipTA, ttgTA, and vanTA transheterozygous flies were reared on food containing a low dose (0.5-10 μg/mL) or a high dose (5-100 μg/mL) of ligand (indicated on the top left-hand side of the plot), no significant reduction of luciferase activity was measured. Only tTA, the positive control, showed a concentration-dependent reduction of luciferase activity. Each dot represents one biological replicate composed of the average of 5 sub-replicates. Exceptions include vanTA and pipTA, where one sub replicate (out of five sub replicates) in one of the high dose treatments (out of 3 replicates) could not be collected due to the difficulty of obtaining sufficient number of transheterozygous flies on their treatments. N represents the total number of flies tested. Bars represent the standard error of the mean (SEM). Significance was determined using a student’s t test. \*\*\*\**p* < 0.0001; \*\*\**p* < 0.0002; \*\**p* < 0.0005; n.s. not significant.

### Attempts to use small molecule ligands to control binary systems

Similar to the Tet-OFF system repressor, which interacts with tetracycline and doxycycline, the repressors from the cymTA, pipTA, ttgTA, and vanTA systems interact with their own specific ligands, corresponding to cumate, virginiamycin M1, phloretin, and vanillic acid, respectively (Mullick et al. 2006; Fussenegger et al. 2000; Gitzinger et al. 2009, 2012)(**Fig. S1B**). Leveraging this prior work, we hypothesized that when no ligand is present, the transactivator should bind to its operon, promoting spatial expression of the reporter genes in the flight muscles (**Fig. 2**). However, when ligand is present and bound to the transactivator, this should result in a confirmational change and temporally prevent the transactivator from binding to its operator. Therefore, the absence of ligand, termed the OFF configuration, should enable the measurement of the maximum spatial gene expression levels of these systems, while the presence of ligand should temporally reduce expression (**Fig. 3B**). To assess this potential, we measured the ability of a small-molecule ligand to repress gene expression in each system. We used three initial ligand concentrations, low dose (0.5-10μg/mL), high dose (5-100 μg/mL), and very high dose (50-1,000μg/mL), though the very high dosage of ligand proved too toxic for fly survival and was excluded. Transheterozygous tTA flies reared on a low dose (1μg/mL) and high dose (10μg/mL) of doxycycline showed a concentration-dependent decrease in GFP fluorescence and firefly luciferase expression (**Fig. 2B** and **Fig. 3B**) (*p* ≤ 0.0005 and *p* ≤ 0.0002, respectively). However, we did not detect a concentration-dependent decrease in luciferase expression in cymTA, pipTA, ttgTA, and vanTA flies (**Fig. 3B**). Confirming this lack of system repression in the presence of the ligand, our fluorescence images indicated that the GFP levels also remained constant for these systems (**Fig. 2B**). This suggests that the highest testable concentrations for each compound (10x) were not sufficient to suppress the cymTA, pipTA, ttgTA, and vanTA systems and may reflect a pharmacokinetics issue of the ligands not reaching the indirect flight muscle and would be worth testing these systems in other tissues *in vivo* in the future.

## Discussion

In this study, we evaluate four bacterially derived transactivation systems *in vivo* in *Drosophila melanogaster*. The use of transgenic binary systems to temporally and spatially control gene expression is one of the most powerful tools in synthetic molecular biology, and these systems assist researchers in modifying cellular functions, generating cellular responses to environmental stimuli, and influencing cellular development (Lewandoski 2001). While established spatial-temporal control systems like GAL4-UAS, Tet-OFF, and the Q-system exist, generating additional systems for the *Drosophila* tool box will be crucial for selectively choosing systems for desired functions or applications. In our work, we demonstrate cymTA, pipTA, ttgTA, and vanTA each robustly and spatially transactivate gene expression in fruit flies, providing new binary transactivation systems that can be used for various research applications.

Using our quantitative luciferase assay, each system was able to demonstrate a higher level of transactivation when compared to the TtA system. This result is promising because higher induction expression systems can be used in several fruit fly applications. Robust expression is generally a desired feature in the development of binary systems. While the tested systems proved to be stronger than TET-OFF, there are still expression level differences between them. For example, vanTA demonstrated the highest average RLU (~360), with both cymTA and pipTa averaging around ~150 RLU, and ttgTA having the lowest average (~75). These differences in gene expression enable the user to choose their desired expression level output (low to high). Even without system repression using their corresponding ligands, these systems still function well for binary gene transactivation.

Since these systems are highly efficient in cell culture (Gitzinger et al. 2009, 2012; Mullick et al. 2006; Fussenegger et al. 2000), they should still be capable of small-molecule control *in vivo,* despite the lack of control we observed in the flight muscle. The Tet-OFF (tTA) positive control suggested our experimental design was sufficient for the activation and suppression of reporter genes. Therefore, it is possible that the amount of compound fed to the animal was not sufficient to either (1) reach the indirect flight muscle tissues or (2) fully suppress the system. The first hypothesis was proposed because in order for the compound to reach the flight muscles, it must pass the midgut and travel through the hemolymph and likely through other organs before reaching the target tissue. To test this hypothesis, the transgenes need to be re-designed to enable expression of reporter genes in the midgut or another easily accessible tissue, where the ligand could more easily reach its target transactivator. Cell culture studies suggest these ligands are able to cross the cell membrane, which indicates these ligands should also be capable of entering animal cells *in vivo*. Whether the ligand is metabolized by the insect, however, is unknown, though could explain the lack of suppression. A comprehensive pharmacokinetics assay may resolve these unknowns.

Our second hypothesis for the lack of ligand repression is that we were unable to reach a concentration that would fully repress the system. In tTA, doxycycline at the 10x concentration was able to fully suppress the system in flies, which may be due to the comparatively lower gene activation level of this system. The concentration of doxycycline tested was sufficient to prevent the transactivator from binding to the operator sequence. Due to the higher levels of transactivation observed in the cymTA, pipTA, vanTA, and to some extent ttgTA systems, it is probable that higher ligand concentrations will be needed for suppression. Because a higher concentration via oral feeding was highly toxic to flies, it would be difficult to conduct such an experiment in a live animal model. Higher concentrations may be achieved via thoracic injection, however, this may be impractical for experiments where tissues are difficult to reach or may otherwise kill the animal. Taken together, we conclusively demonstrated that these bacterially-derived systems can robustly function as genetic binary transactivational systems *in vivo* and these tools expand the molecular genetic *Drosophila* toolbox. Our work provides the first step in the characterization of new transactivation systems in fruit flies and may contribute to the generation of novel synthetic tools that can be used in other animal systems, perhaps even mosquitoes (Zhao, Tian, and McBride 2020) to elucidate molecular genetic questions and to design advanced biological circuits.

## Material and Methods

### Plasmid construction

For the construction of driver transgenes, the following cloning strategies were first performed: To generate the vanTA driver (vector 907F1), an attB cutter backbone containing a multiple cloning site was digested using restriction enzymes SwaI and XbaI. The following PCR products were inserted via Gibson cloning: a Flightin (*Fln*) promoter fragment amplified from *Drosophila* genomic DNA using primers 907F1.F1 and 907F1.R1, a vanR repressor protein from a gene-synthesized plasmid using primers 907F1.F2 and 907F1.R2, and finally, a VP16-SV40 fragment amplified from a gene-synthesized plasmid using primers 907F1.F3 and 907F1.R3. To generate the cymTA driver (vector 907H1), an attB cutter backbone containing a multiple cloning site was digested using restriction enzymes SwaI and XbaI. The following PCR products were inserted via Gibson cloning: a Flightin (*Fln*) promoter fragment amplified from *Drosophila* genomic DNA using primers 907F1.F1 and 907H1.R1, a cymR repressor protein from a gene-synthesized plasmid using primers 907H1.F2 and 907H1.R2, and finally, a VP16-SV40 fragment amplified from a gene-synthesized plasmid using primers 907H1.F3 and 907F1.R3.

In a separate cloning strategy, we cloned in a longer variant of the VP16 domain (VP16’) in the ttgTA and pipTA systems to see if this VP16’ would also provide robust activation of reporter genes in our system. We used vector 907F1 as a backbone to save two PCR amplification steps (Fln promoter fragment and the SV40 fragment). To generate the pipTA driver (vector 907K), we digested plasmid 907H1 with AscI and BglII restriction enzymes. The following PCR products were inserted via Gibson cloning: the pipR repressor sequence amplified from a gene-synthesized plasmid using primers 907K.F2 and 907K.R2, and the VP16 sequence amplified from a gene-synthesized vector with primers 907K.F3 and 907K.R3. To generate the ttgTA driver (vector 907L), we digested plasmid 907H1 with AscI and BglII restriction enzymes. The following PCR products were inserted via Gibson cloning: the ttgR sequence amplified from a gene synthesized plasmid using primers 907L.F2 and 907L.R2, and the VP16 sequence which was amplified from a gene synthesized vector with primers 907L.F3 and 907L.R3. We chose the Tet-OFF system as the positive control to compare novel systems to a widely used repressible system. For this, the TetR-VP16 sequence was amplified from an Oxitec plasmid OX1124 (Morrison et al. 2012) using primers 1025.c1 and 1025.c2 and cloned into a AscI/BglII digested 907F1 vector. All primers used for driver constructs are listed in **Table S1**.

To engineer responder transgenes, several cloning steps were performed. First an attB cutter vector was digested with AscI and XbaI. The following were added via Gibson cloning to create intermediate vector 908-1a: firefly luciferase, amplified from a gene synthesized vector using primers Firefly.F and Firefly.R and a T2A-GFP-p10-3’UTR sequence amplified from a previously described vector (Kandul et al. 2019) using primers GFP.F and GFP.R. Then, 908-1a was digested with XhoI, and the following components were added via Gibson cloning to generate intermediate vector 908-1b: an Hsp70 minimal promoter, amplified synthetically using primers Hsp70.F and Hsp70.R and a UASp promoter (Rørth 1998) amplified from the pWALIUM22 plasmid using primers UAS.F and UAS.R. Finally, 908-1b was digested with XbaI to add an SV40-Renilla-luciferase-ubiquitin sequence that was amplified from a previously engineered vector 1052 (unpublished) with primers UbiqRen.F and UbiqRen.R to make vector 908-1c. Then, OA1c was digested with XhoI/PacI to insert the operator sequences for each system upstream of the minimal Hsp70 promoter. Specifically, the operator sequences (ttgO) for the ttgTA system were amplified from a gene-synthesized vector using primers ttgO.F and ttgO.R to make the final vector 908E.

For the pipTA system, operator sequences (pipO) were amplified synthetically using primers 908A17 and 908A18 to make the final vector 908G. For the tTA system, operator sequences (tetO) were synthetically amplified using primers 908A11 and 908A12 to make the final vector 908H. For the vanTA system, the operator sequences (vanO) were amplified using primers 908A13 and 908A14 to make the final vector 908I. For the cymTA system, operator sequences (cymO) were amplified using primers 908A15 and 908A16 to make the final vector 908J. All primers used for responder constructs are listed in **Table S2**. For a complete list of vectors, Addgene ID numbers, and vector descriptions, please refer to **Table S3**. Plasmid DNA and complete annotated DNA plasmid sequences maps can be found at www.Addgene.com.

### Fly rearing and genetic crosses

Rainbow Transgenics (Camarillo, CA, USA) performed all of the fly injections. All driver and responder constructs were injected into a transgenic 3^rd^ chromosome attP site line marked with 3xP3-RFP (Bloomington Drosophila Stock Center (BSC), Bloomington, IN, USA; RRID: BDSC_24486, y[1] M{RFP[3xP3.PB] GFP[E.3xP3]=vas-int.Dm}ZH-2A w[*]; M{3xP3-RFP.attP’}ZH-86Fa). Recovered transgenic lines containing the construct at the 3^rd^ chromosome site were balanced on the 3^rd^ chromosome using a double-chromosome balancer line (w1118; CyO/Sp; Dr/TM6C, Sb1). Single homozygous transgenic driver and responder flies were maintained as separate lines. Flies were maintained on cornmeal, molasses, and yeast medium (Old Bloomington Molasses Recipe) at 25°C with a 12H/12H light/dark cycle. To assess system activation, we used Instant Drosophila Food (Formula 4–24) from the Carolina Biological Supply Company. In each fly vial (FlyStaff.com), 1.1 g of dry food was mixed with 5 ml of distilled water. To obtain transheterozygous flies, driver strains were crossed to the responder strains in treated or non-treated food. As a control, the responder strain was crossed to a WT (w[1118]) strain to produce heterozygous responders. To assess system suppression with ligand, instant food was supplemented with doxycycline, cumate, phloretin, vanillic acid, or virginiamycin M1 in varying concentrations using the compound solutions described below. All driver and responder strains were deposited to the Bloomington Drosophila Stock Center, and their corresponding BDSC IDs are listed in Table S3.

### Compound solutions

Doxycycline (195044, MP Biomedicals LLC), with a half-life of 11–12 hrs (Graham and Pile 2016), was prepared as a stock solution of 1,000 μg/mL in 100% ethanol. To make concentrations of 1 μg/mL and 10 μg/mL for 1x and 10x treatments, respectively, stock solution was diluted in distilled H_2_O. Cumate (QM100A-1, SBI) arrived as a 300 mg/mL (10,000X) stock solution in 500 μL. To make 1x and 10x treatment solutions, two serial dilutions at 1:10 were first performed with distilled H_2_O to reach a workable concentration of 300 μg/mL. Then 2.5 μg/mL (low treatment) and 25 μg/mL (high treatment) working concentrations were generated in distilled H_2_O. Phloretin (P7912, Sigma-Aldrich), with a half-life of 70 hrs (Gitzinger et al. 2009), was prepared as a stock solution of 10 mg/mL in 100% ethanol. The phloretin stock solution was diluted in distilled H_2_O to 4 μg/mL and 40 μg/mL for the low and high ligand treatments, respectively. Vanillic acid (H36001, Sigma-Aldrich), with a 7 min half-life (Yrbas et al. 2015), was prepared as a stock solution at 1,680 μg/mL by dissolving the powder in distilled H_2_O over a hot magnetic spin plate. Since vanillic acid is an acidic compound, 3 M of NaOH was added to neutralize the solution to a pH of 7.0 for fly food. Working solutions were made by diluting the stock solution in distilled H_2_O to a final concentration of 10 μg/mL and 100 μg/mL for the low and high ligand treatments, respectively. Virginiamycin M1 (V2753, Sigma-Aldrich), with a half-life of 4–5 hrs (Gitzinger et al. 2009; Kwon 2017), was prepared as a stock solution at 500 μg/mL in 100% ethanol. Working concentrations of 0.5 μg/mL and 5 μg/mL for the low and high ligand treatments respectively, were made by diluting the stock solution in water. Fly food treatments were set up by adding 1.1g of Formula 2-24 Instant Drosophila Medium (#173218, Carolina) to an empty fly vial and adding 5mL of working solution. The solution was allowed to sit for at least 4 hours before adding flies.

### Imaging

Flies were scored and imaged on the Leica M165FC fluorescent stereomicroscope equipped with the Leica DMC2900 camera. Images were done under constant conditions.

### Luciferase assays

The Dual-Luciferase® Reporter Assay System (Promega, Madison, WI) was used to measure firefly luciferase expression in response to transactivation or repression. To measure luciferase consistently, 3–day-old male flies were individually collected in a 1.5 μL microcentrifuge tube and stored at −80°C before lysing. Treatments were done in triplicate. Each replicate contained five sub-replicates with 5 flies each. Therefore, a total of 25 flies were tested per replicate. To lyse the sample, the Passive Lysis Buffer 5X from the Dual-Luciferase® Reporter Assay System kit (Promega, Cat.# E1910) was diluted in distilled H_2_O at 1:5 to make a 1x lysis buffer. Then, 40 μL of lysis buffer was used to mechanically disintegrate the sample with a plastic pestle, and an additional 40 μL of buffer was used to wash the remaining tissue off the pestle into the tube. To remove the tissue, lysed samples were subjected to a 15 min centrifuge spin at 10,000 rpm. Then, 75 μL of the supernatant (without tissue) was removed and placed into a clean tube and stored at −80°C. Before measuring luciferase activity, Luciferase Assay Reagent II (LARII) and Stop and Glo® reagents from the same Dual-Luciferase® assay kit were prepared according to the manufacturer’s instructions. Firefly luciferase activity was first measured by adding 100 μL of LARII in a tube containing 5 μL of lysed sample, which was then placed in the GloMax® 20/20 Luminometer (Promega, Madison, WI) to measure luminescence in relative luciferase units (RLU) for an integration time of 10 s. Then, 100 μL of Stop and Glo® was added to measure Renilla luciferase for 10 s. Each measurement was recorded in an Excel spreadsheet and later organized for calculations.

### Normalization of luciferase and statistical methods

The quantitative results are expressed as relative luminescence units (RLU) and are normalized by taking the ratios of firefly/Renilla luciferase. To determine the relative luciferase activity (RLA), firefly/Renilla ratios were first calculated for each individual sample from acquired luminometer data. Then the following formula from (Potter et al. 2010) was used to calculate RLA:

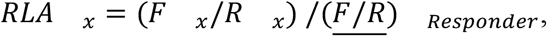

where

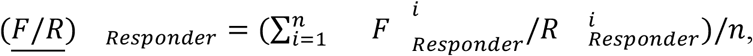

Where n = number of responder-only samples; F = firefly luciferase RLU; R = Renilla luciferase RLU. The average and SEM were determined for each treatment, and the statistical significance was determined using a Student’s t test. Comparisons were considered statistically significant with *p* < 0.05. GraphPad Prism version 8.3.1 for macOS was used for these analyses (GraphPad Software, San Diego).

### Transactivator Modelling

The tertiary structure of the binary proteins was modeled using LOMET online, a metaserver-based protein fold recognition program (Zheng et al. 2019; Wu and Zhang 2007). With this online tool, 3D models were generated by collecting high-scoring structural templates that were compared with the crystal structure of repressor proteins characterized in the literature.

## Author Contributions

O.S.A conceived and designed experiments. S.G. obtained genetic cross data and analyzed all the data. L.C.V and S.C.M performed molecular analyses. All authors contributed to writing and approved the final manuscript.

## Acknowledgments

This work was supported in part by UCSD lab startup funding and a National Institutes of Health New Innovator Award (1DP2AI152071-01) awarded to O.S.A.

## Ethical conduct of research

We have complied with all relevant ethical regulations for animal testing and research and conformed to the UCSD institutionally approved biological use authorization protocol (BUA #R2401).

---

**Figure S1.**
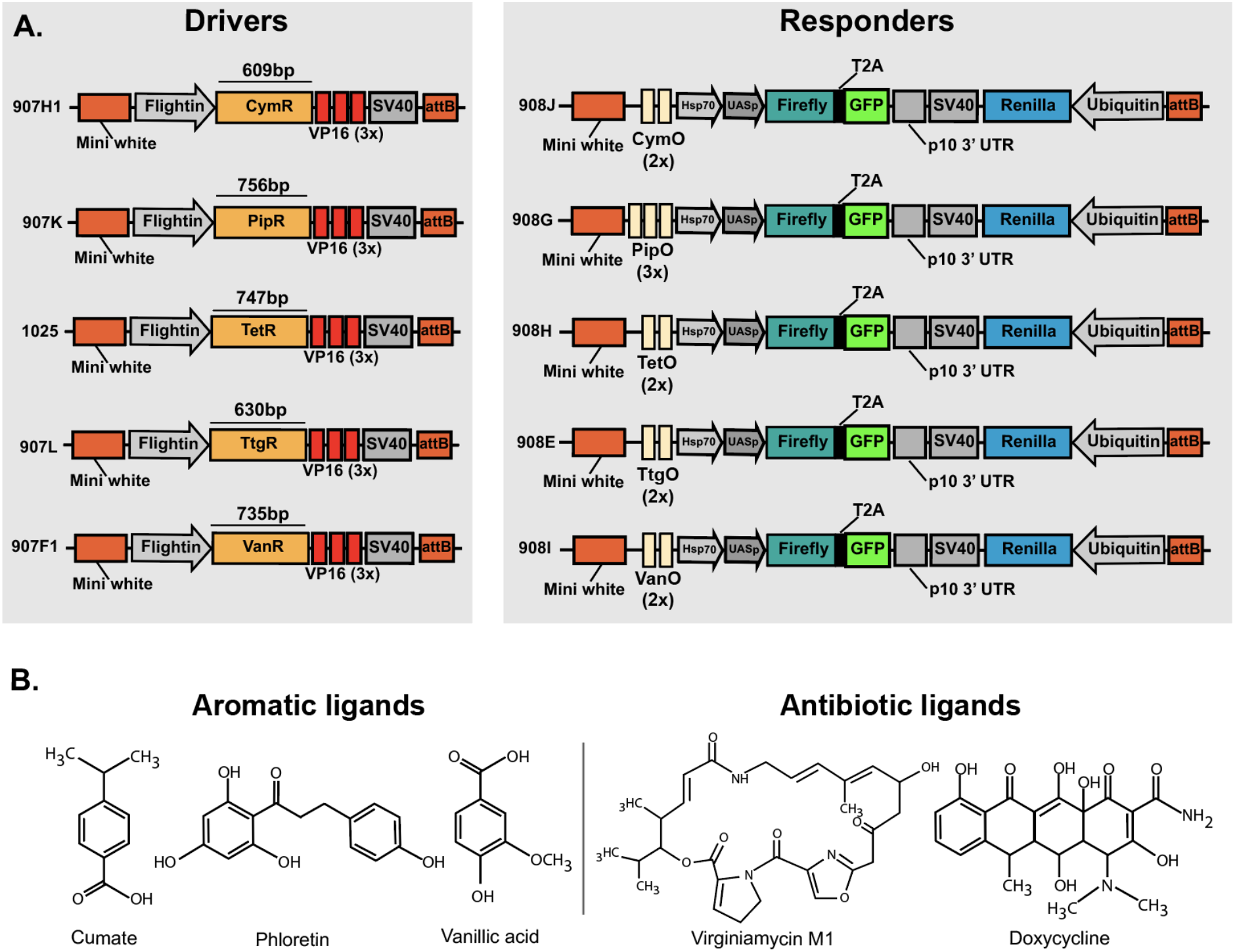
Schematic representation of driver and responder transgenes tested for each system. (A) Each regulation system described in this study is distinguished by its repressor sequence, operon sequence, and ligand used to suppress the system. To compare all these systems in *Drosophila melanogaster*, the rest of the components were kept consistent between transgenes. (B) Chemical structures of ligands tested in this study. These ligands can be characterized as aromatic or antibiotic.

**Table S1.**
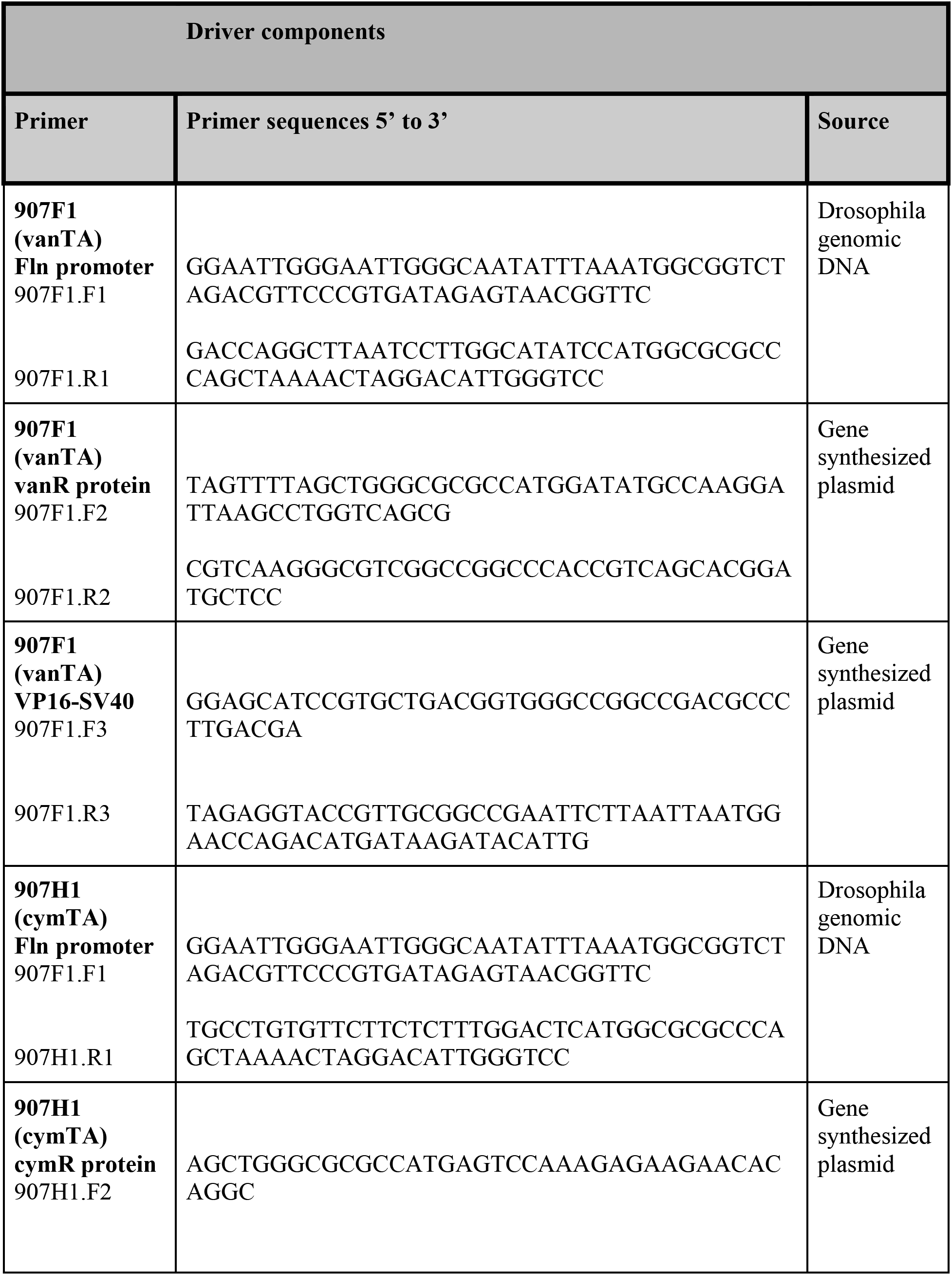
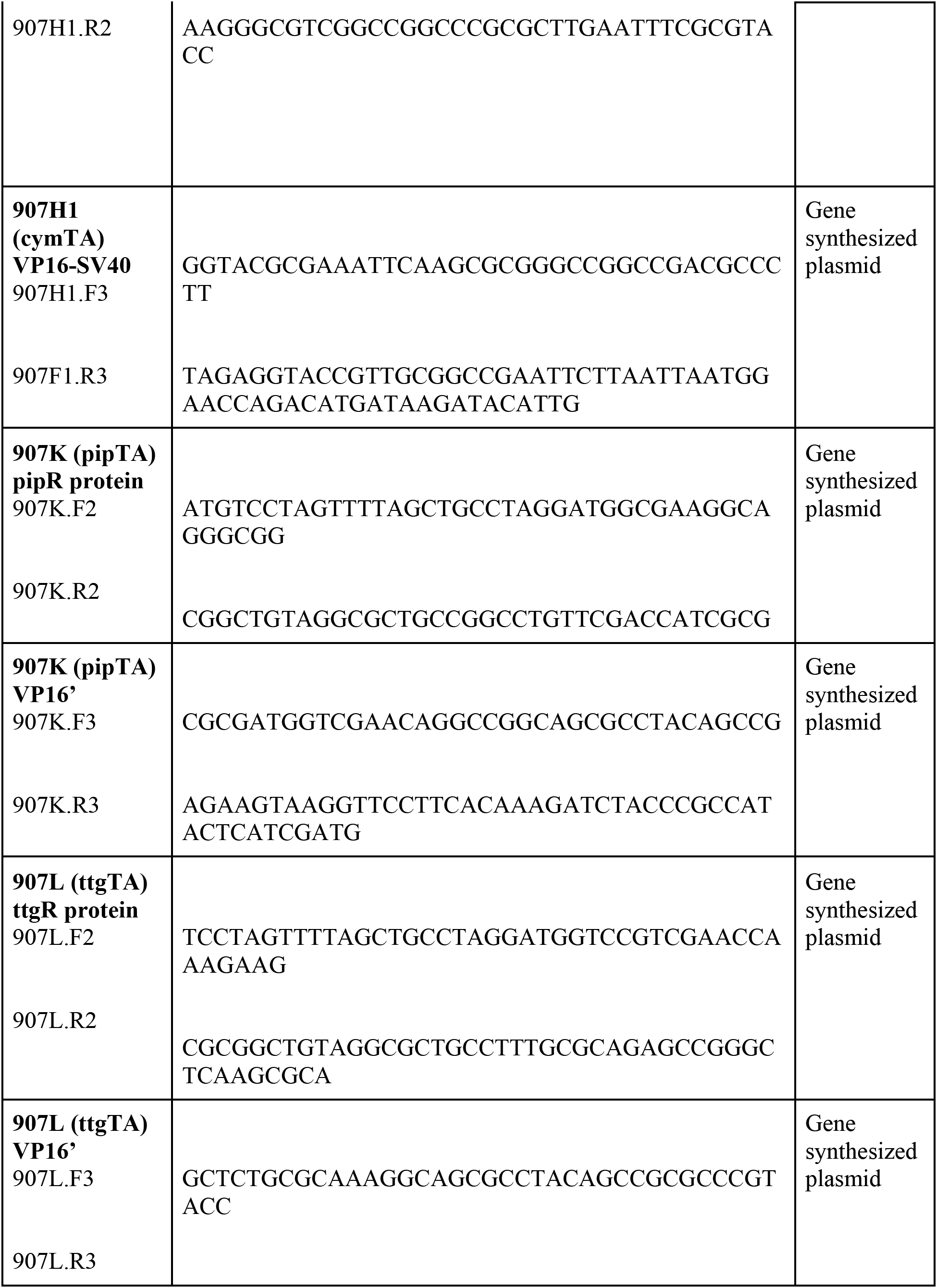
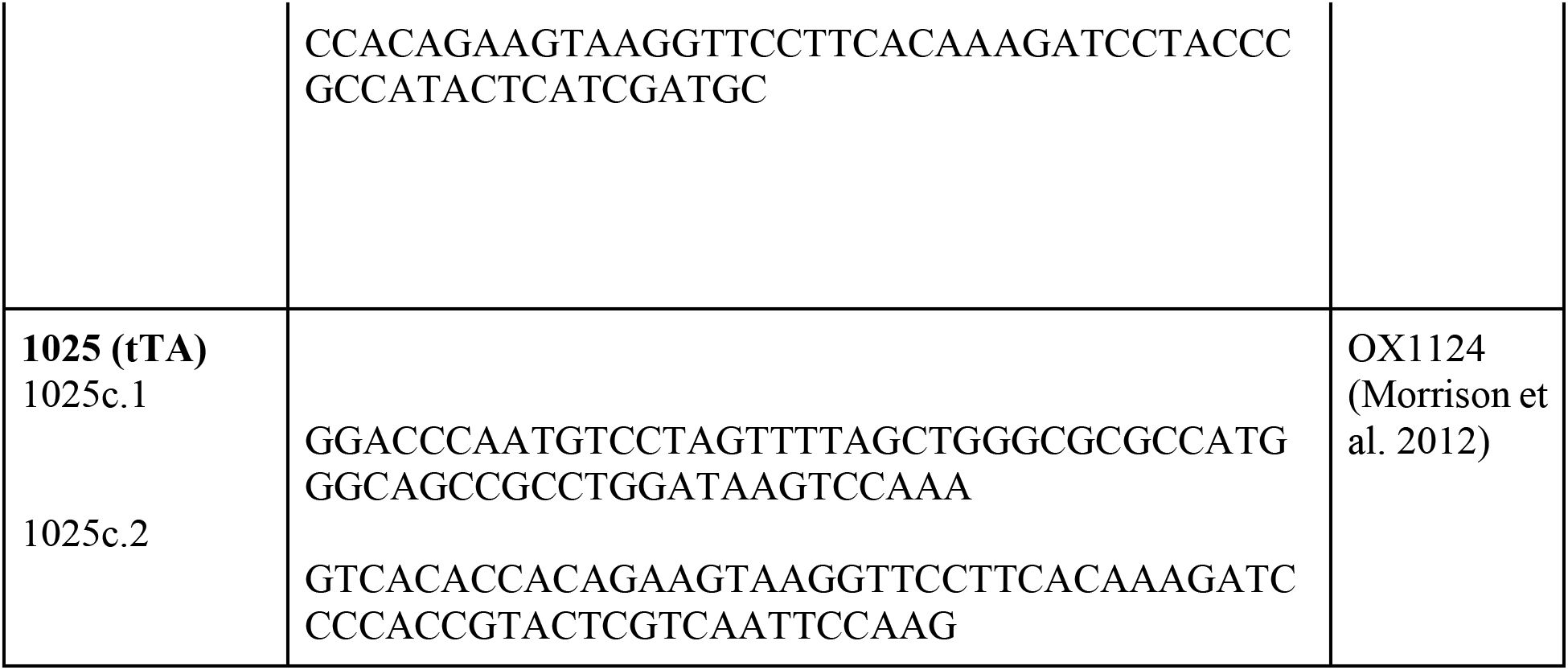
Primers used to build synthetic driver constructs.

**Table S2.**
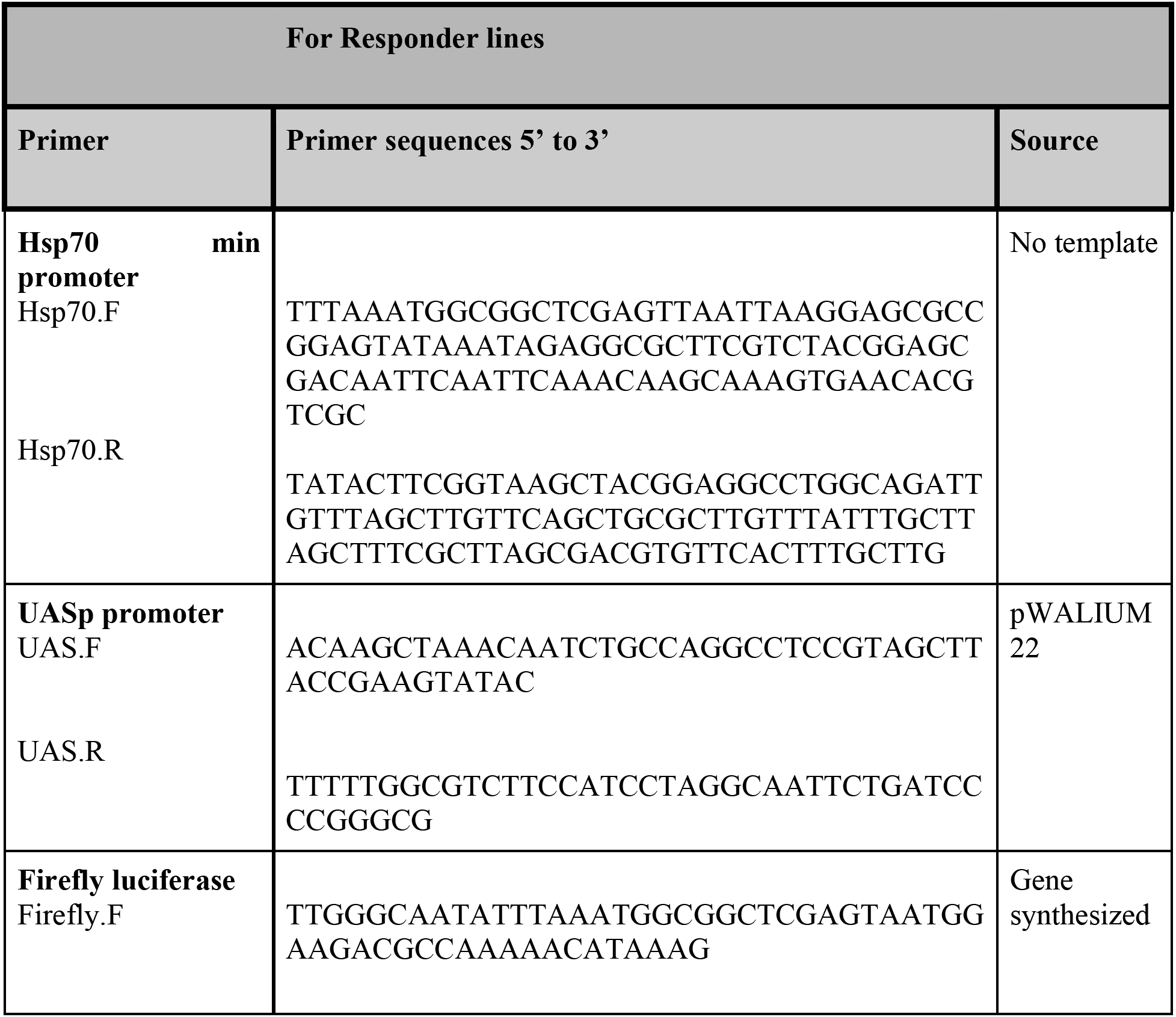
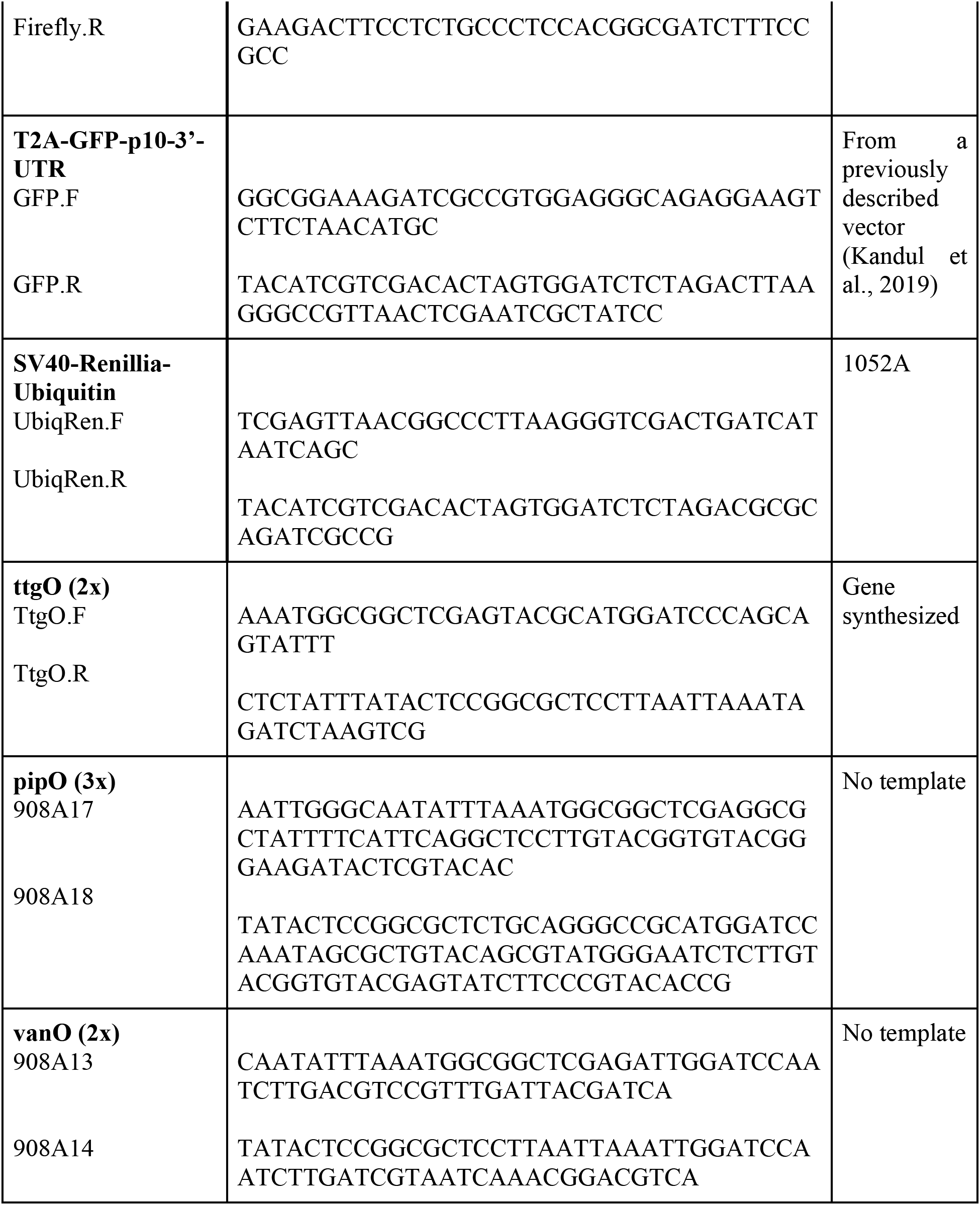
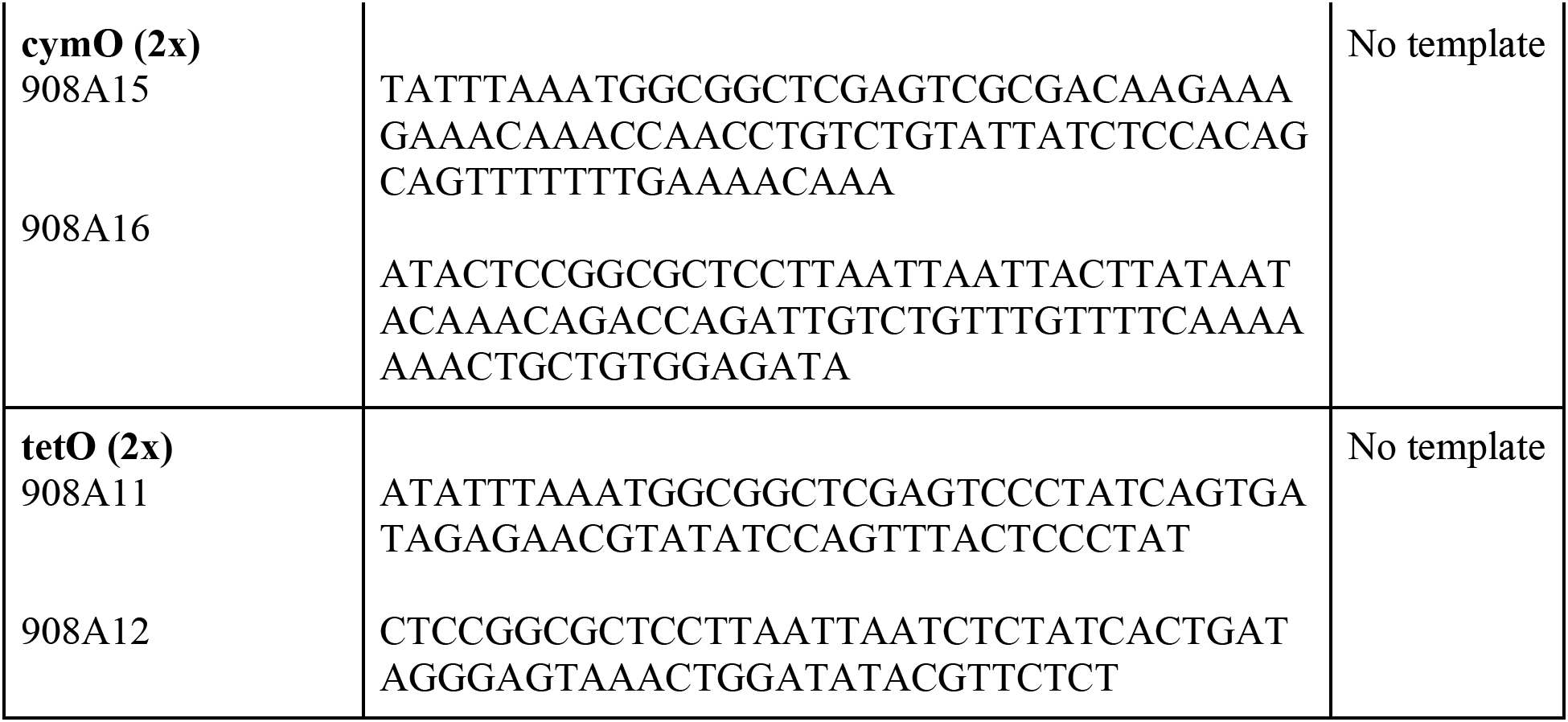
Primers used to build responder constructs.

**Table S3.** Final vector constructs and their descriptions. List of vectors and transgenic fly lines generated in this study.

| Vector # | Addgene ID | BSC ID | FlyBase symbols | Description |
| --- | --- | --- | --- | --- |
| **907F1** | 160233 | 91379 | w[*];M{RFP[3xP3.PB]<br>w[+mC]=fln-vanTA.G}ZH-86Fa | mini-w+-FlightN-vanTA-SV40-attB |
| **907H1** | 160234 | TBD | M{fln-cymTA.G}ZH-86Fa | mini-w+-FlightN-cymTA-SV40-attB |
| **1025** | 160235 | 91381 | w[*];M{RFP[3xP3.PB]<br>w[+mC]=fln-tTA.G}ZH-86Fa | mini-w+FlightN-tTA-SV40-attB |
| **907K** | 160236 | 91380 | w[*];M{RFP[3xP3.PB]<br>w[+mC]=fln-pipTA.G}ZH-86Fa | mini-w+-FlightN-pipTA-SV40-attB |
| **907L** | 160237 | TBD | M{fln-ttgTA.G}ZH-86Fa | mini-w+-FlightN-ttgTA-SV40-attB |
| **908E** | 160238 | 91382 | w[*];M{RFP[3xP3.PB]<br>w[+mC]=ttgO-UASp-<br>pLUC.T2A.EGFP,Ubi-<br>rLUC}ZH-86Fa | mini-w+-ttgO(2x)-Hsp70min-UASpmin-<br>Firefly-T2A-eGFP-p10-3'UTR-SV40-<br>Ubiquitin-Renilla-attB |
| **908G** | 160239 | 91383 | w[*];M{RFP[3xP3.PB]<br>w[+mC]=pipO-UASp-<br>pLUC.T2A.EGFP,Ubi-<br>rLUC}ZH-86Fa | mini-w+-pipO(3x)-Hsp70min-UASpmin-<br>Firefly-T2A-eGFP-p10-3'UTR-SV40-<br>Ubiquitin-Renilla-attB |
| **908H** | 160240 | 91384 | w[*];M{RFP[3xP3.PB]<br>w[+mC]=tetO-UASp-<br>pLUC.T2A.EGFP,Ubi-<br>rLUC}ZH-86Fa | mini-w+-tetO(2x)-Hsp70min-UASpmin-<br>Firefly-T2A-eGFP-p10-3'UTR-<br>Ubiquitin-Renilla-attB |
| **908I** | 160241 | 91385 | w[*];M{RFP[3xP3.PB]<br>w[+mC]=vanO-UASp-<br>pLUC.T2A.EGFP,Ubi-<br>rLUC}ZH-86Fa | mini-w+-vanO(2x)-Hsp70min-UASpmin-<br>Firefly-T2A-eGFP-p10-3'UTR-SV40-<br>Ubiquitin-Renilla-attB |
| **908J** | 160242 | TBD | M{cymO-UASp-<br>pLUC.T2A.EGFP,Ubi-<br>rLUC}ZH-86Fa | mini-w+-cymO(2x)-Hsp70min-<br>UASpmin-Firefly-T2A-eGFP-p10-<br>3'UTR-SV40-Ubiquitin-Renilla-attB |

